# Comparison of co-immunoprecipitation techniques for effective identification of SNTA1 interacting proteins in breast cancer cells

**DOI:** 10.1101/2022.03.16.484629

**Authors:** Saima Sajood, T. S. Keshava Prasad, Basharat Bhat, Zuhiab F. Bhat, Riaz A Shah, Hina F Bhat

## Abstract

Diverse signal transduction pathways involve thousands of proteins acting in concert that contribute to breast cancer pathology. SNTA1 has emerged as a common link that functions to wire various complex signaling circuits within the cell. In our study we used endogenous SNTA1 protein as bait in pull-down assays to identify SNTA1 interactome from breast cancer cell extract. We compared the utility of two co-immunoprecipitation techniques conventional co-IP and cross-linking co-IP coupled to mass spectrometry (IP–MS) that intriguingly generated valuable inventory of SNTA1 associated proteins in breast cancer cells that may or may not interact physically but contribute to shared functions in breast cancer pathology. We observed that Pierce cross-link IP kit enhanced sensivity of the analysis as it decreased the number of background proteins with a remarkable increase in identified list of proteins. We have also observed that different approaches are able to consistently detect the same signaling outcome as observed by Gene Ontology. Identification of interacting partners of SNTA1 in breast cancer cell lines may depict the possible signaling mechanism of this protein in molecular pathogenesis of breast cancer and its possible use as a therapeutic target.

## Introduction

Breast cancer has surpassed lung cancer as the most commonly diagnosed cancer among women worldwide, with an estimated 2.3 million new cases (11.7% of total cases) in 2020. Despite extensive research, breast cancer is still fifth leading cause of cancer-related deaths worldwide, with 685,000 deaths (Sung et al., 2021). Thus, more studies are needed to investigate the underlying molecular mechanisms involved in mammary tumor progression and metastasis. α-1 syntrophin (SNTA1) is a distinguishing member of PDZ family with multiple structural domains that facilitate crosstalk between various signaling proteins or cytoskeletal components. The unique domain architecture of PDZ domain within PH domain of SNTA1 builds a platform for formation of signaling complexes with a variety of partners including membrane channels, receptors, kinases and other signaling proteins (Kyle & Makielski 2014; Huang et al., 2020). SNTA1 is able to interact with multiple isoforms of hetero-trimeric G proteins including Gs and Gα-family proteins (Gαq, Gαo, Gαs and Gαi) suggesting its role in signaling pathways via GPCRs. (Akiko, et al., 2008). It provides a link between extracellular components and cytoskeletal/signaling proteins via Dystrophin glycoprotein complex (DGC) (Court et al., 2005; Bhat et al., 2018; Adams et al 2018).

Intriguingly SNTA1 promotes cell proliferation, cell migration and production of reactive oxygen species via Rac1 activation in breast cancer cell lines (Iwata et al., 2004; Bhat et al., 2014). However, SNTA1 binding partners that may modulate or regulate its activity during various pathological conditions have not been fully elucidated.

Network -based analyses combined with omics datasets has enhanced identification of novel cancer-associated genes and underlying molecular mechanisms (Li et al., 2016). Thorough understanding and identification of SNTA1-associated proteins in breast cancer cells may accelerate comprehensive identification of novel biomarkers or targets for disease prognosis and their relevant biological functions. In the present study, we carried out co-immunoprecipitation (co-IP) assays coupled with LC-MS/MS approach which provides a rapid, sensitive, and reliable experimental strategy to identify protein interactors, and has proved to be an alternative for the discovery of novel interacting partners that cannot be detected using yeast-two hybrid analyses. We performed both conventional and crosslinking co-IP methods, however crosslinking outperformed the standard co-IP method which enabled identification of large number of SNTA1 associated (in vivo) direct or indirect interactions or protein complexes.

## Materials and Methods

### Reagents

Dulbecco’s modified Eagle’s medium (DMEM) was the product of HyClone, USA. Trypsin and FBS were purchased from Gibco, USA. Pierce crosslink IP kit was purchased from Thermo scientific. The primary antibody against SNTA1 was purchased from Life Span. Protein A/G plus agarose suspension solution was obtained from Calbiochem (MERCK) and Dimethyl Sulfoxide (DMSO) was obtained from Amresco. 100 X protease inhibitor cocktail, PBS (pH 7.4, 795 mg/L Na2HPO4, 144 mg/L KH2PO4, 9000 mg/ mL NaCl, without calcium and magnesium) and Laemmli buffer (2% SDS, 10% glycerol, 0.002% bromophenol blue, 125 mM Tris HCl) were purchased from Himedia.

### Cell Culture

Human breast cancer cell line MCF-7 was obtained from the NCCS Pune. Cells were maintained in DMEM (Hyclone) with 10% (v/v) fetal bovine serum (FBS), 2 mm l-glutamine, 50 units/ml penicillin G, and 50 μg/ml streptomycin in a CO_2_ incubator at 37°C. After rinsing twice with cold PBS, cells were lysed in NP-40 lysis buffer containing 1% NP-40 and protease inhibitors on ice for 45 min with occasional vortexes. After centrifugation at 15,000 g for 10 min at 4°C, protein was quantified by the protein assay kit (Qubit 2.0). Equal amount of samples was run on a 10% sodium dodecyl sulphate polyarcylamide gel electrophoresis (SDS-PAGE).

### Conventional method using Protein G agarose beads

Protein G coupled agarose beads used for pull down were washed three times with cold PBS and incubated with desired antibody (3 μg) overnight on rotatory shaker (17rpm at 4°c). On the following day, cellular protein lysates collected from different samples were added to the prepared antibody conjugated agarose beads and incubated on rotatory shaker for 4 hours. This was followed by centrifugation at 2000 rpm for two minutes at 4°c. To remove unbound proteins from beads washings were done with ice cold PBS followed by mixing with 2x sample buffer and boiled for 5 min. at 95°C. The eluted protein was either used for LCMS/MS analysis or was electrophoresed on 10% SDS-PAGE for western blotting analysis.

### Pierce Crosslink IP Kit

This technique involves covalent crosslinking of antibodies onto Protein A/G resin which facilitates the more effective and efficient purification of target protein without contamination by the antibody and increases the ability to more effectively wash and separate samples from the beaded agarose resin (**Figure 1**).

**Figure 1:**
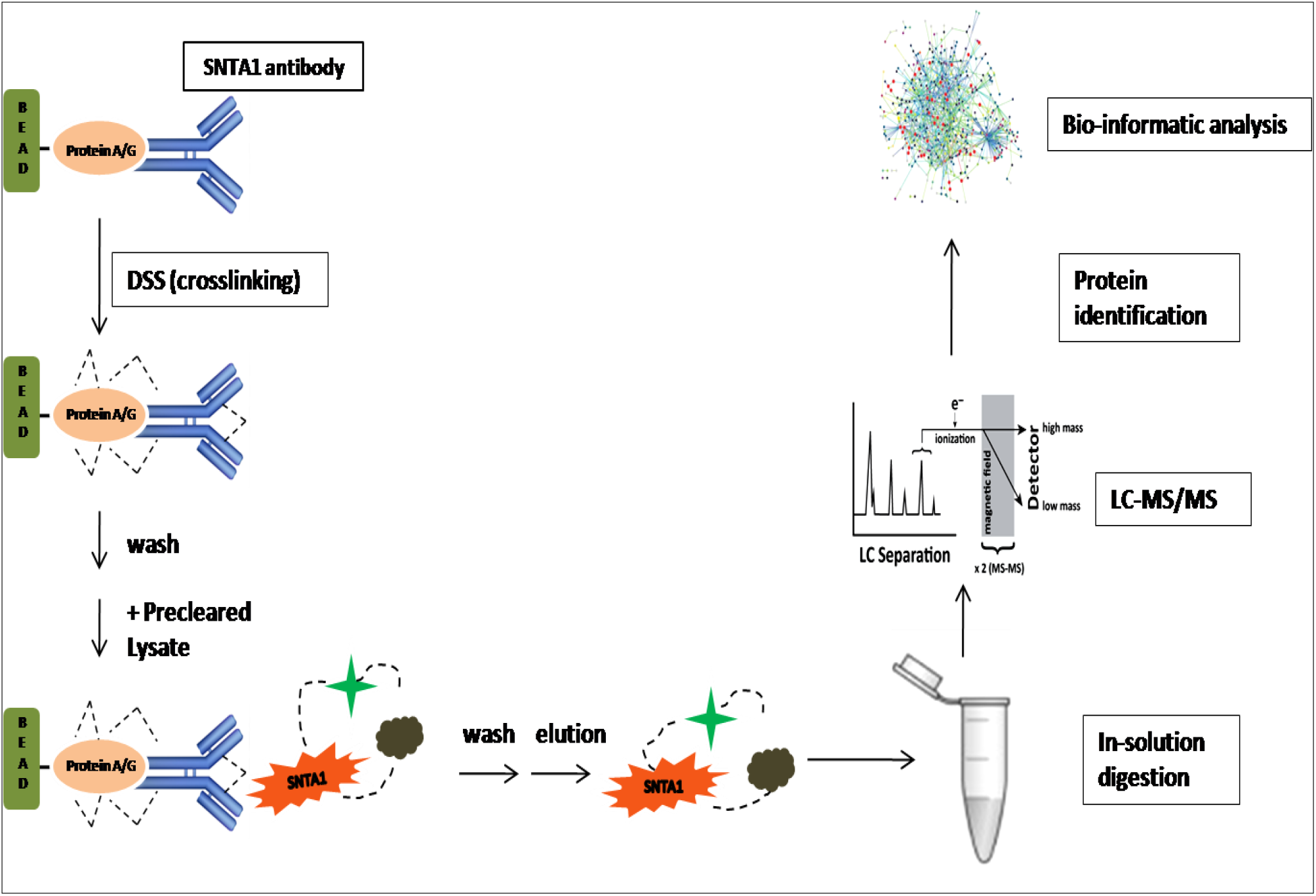
Crosslinking approach to identify SNTA1 interacting proteins.

For Antibody binding to Protein A/G Agarose, resin was first washed with 1x coupling buffer twice and 10µg of desired antibody and 1x coupling buffer were added directly to the resin in the column, volume was adjusted to 100µl with milli-Q water. The slurry was incubated on rotatory shaker overnight at room temperature. Following day, the resin was washed with coupling buffer twice and flow-through was discarded. Further crosslinking of the bound antibody was done to diminish co-elution of antibody with antigen during the elution steps. 2.5µl of 20xcoupling buffer, 9µl of of 2.5 mM of DSS (disuccinimidyl suberate) and 38.5 µl of Milli-Q were added to the column such that total volume was adjusted to 50µl.

After incubating the reaction mixture for 30-60 mins at RT, non-crosslinked antibody was removed and crosslinking reaction was quenched using elution buffer followed by washing of column with cold IP lysis/wash Buffer.

Before proceeding for immunoprecipitation protocol, protein cell lysates were cleared using 80 µl of the control Agarose Resin slurry per mg of lysate in spin column and incubated at 4°c for 60min with gentle end-over-end mixing. Centrifugation was done at 1000g for 1 minute and flow through was added to immobilized antibody. Pre-cleared lysate was incubated/mixed with antibody-crosslinked resin in the column for 1-2 hours or overnight at 4°c. Washing of the column was done with 200µl IP Lysis/wash buffer twice and single wash with 1X conditioning buffer. Final elution was done in 50µl elution Buffer and eluate was analyzed.

### In-solution digestion

The pool of immune-precipitated protein samples were first reduced using DTT (5mM) at 60°C for 45 min and then alkylated using Iodoacetamide (IAA) at RT in dark for 10 minutes. After this, proteins were enzymatically digested using sequencing grade modified trypsin (Promega) overnight at 37ºC. Obtained peptide extracts were vacuum dried and C_18_ clean up was carried out.

### LC-MS/MS analysis

Cleaning of Peptide extracts was performed using C18 and SCX StageTip based method for C18 solvent-A (0.1% formic acid) for equilibrating column and elution was done in solvent (40% Acetonitrile and 0.1% formic acid) and then vacuum dried for SCX 0.1% trifluroactic acid was used for equilibrating column 0.2% trifluroactic acid for cleaning elution was done with 500mM of ammonium acetate in 50% on acetonitrile. The peptides were resuspended in formic acid (0.1%), loaded onto a 2 cm trap column and were separated using a 15 cm analytical column at a flow rate of 300 nl/min for 120 minutes/fraction. Orbitrap mass analyzer was used for global MS survey scan performed at a scan range of 400-1600 m/z mass range in a data-dependent mode for a maximum injection time of 5 minutes.

Data obtained for MS/MS analysis at top at speed mode with 3s cycles was further subjected to higher collision energy dissociation with 34% normalized collision energy. Orbitrap mass analyzer was used to perform MS/MS scans and internal calibration was done using lock mass option from ambient air.

## Results

### Conventional coimmunoprecipitation

Co-immunoprecipitation is considered as standard approach for identification of large scale protein–protein interactions in vivo. Conventional Co-IP experiments can detect various proteins that interact directly or indirectly to form protein complexes. Here, we used SNTA1 specific antibody in conjunction with Protein A/G affinity beads to identify SNTA1 interacting proteins present in cell lysates. Experiments were performed using whole cell lysates, without any conscious overexpression of alpha 1 syntrophin protein so that relative concentrations of protein components remain unchanged. Total protein components isolated using this technique were gel extracted, trypsin digested and analyzed by liquid chromatography tandem mass spectroscopy (LC-MS/ms). The raw data obtained from above analysis was searched against Human refseq 81 database using Proteome Discoverer software suite version 2.2 (Thermo Fisher Scientific). Protein sequences were obtained from NCBI, and SEQUEST & Mascot algorithms were used for searching MS/MS data against the protein database along with known mass spectrometer contaminates.

To distinguish between the bonafide SNTA1 binding partners and non specifically bound background proteins, we used serum IgG antibody of same concentration, as that of SNTA1 antibody, as negative control in the study. We derived a total of 17 proteins which were compared to the negative control to filter out common proteins or background proteins. Common proteins identified ACTA1, HNRNPA2, TUBA1C, Nucleolin, Stress-70 protein and Filamin A isoform 2 in both the samples were considered to be background proteins. Thus using this approach we identified 11 SNTA1 binding partners which include HIST1H2B, HSPB1, AHNAK, HSP90 ALPHA isoform I, Per oxiredoxin-4 isoform X1, Protein S100-A11, Tropomyosin alpha 3 chain isoform, Keratin type I cytoskeletal 18, 14-3-3 protein episilon, E3 ubiquitin –protein ligase RNF213 isoform X2. The majority of the proteins appear to have roles in actin organization and cell-cell adhesion. Cell adhesion molecules significantly contribute to cancer progression and metastases (**Figure 2**).

**Figure 2:**
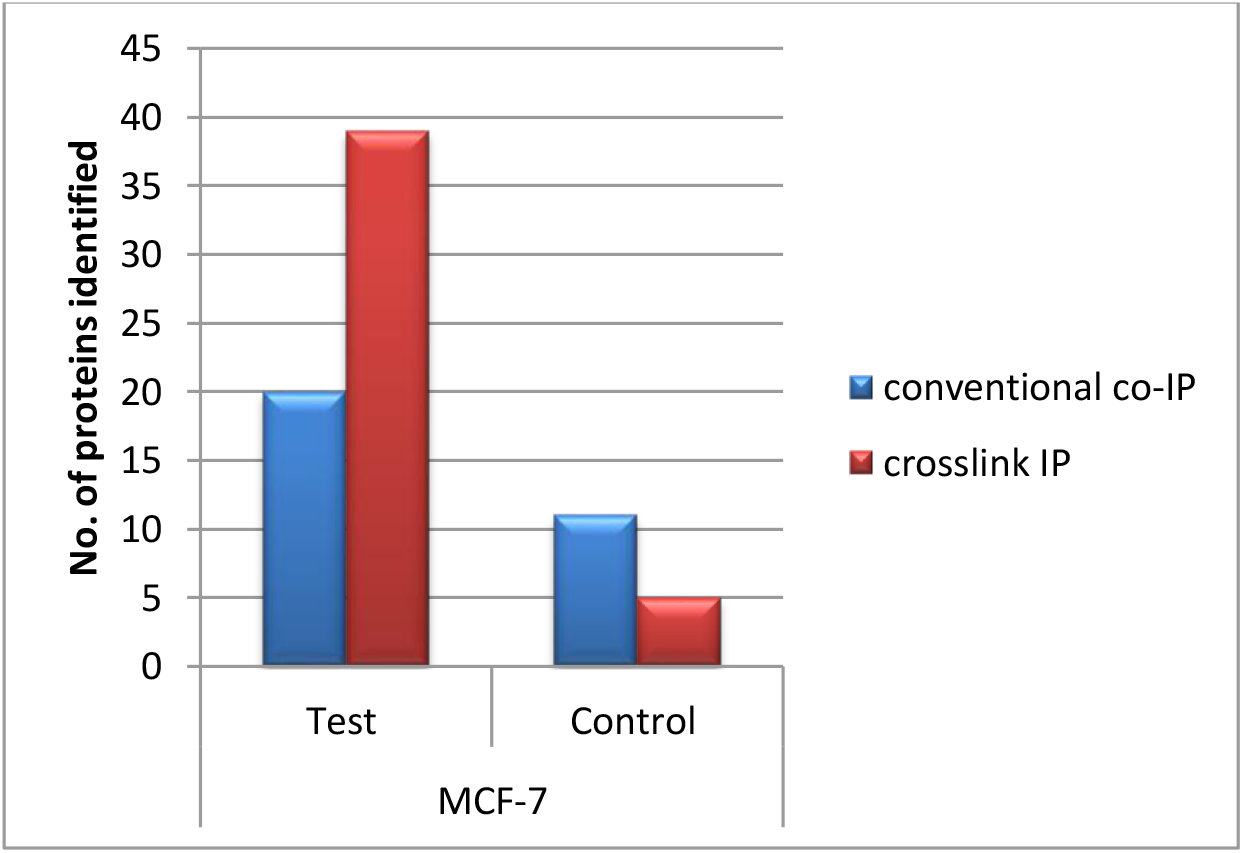
showing efficiency of two approaches (conventional and crosslink IP) for determining SNTA1 interactome in MCF-7 cell line.

### Chemical crosslinking

Chemical crosslinking using DSS (disuccinimidyl suberate) generated abundant SNTA1 interacting proteins with a decrease in background proteins presumably due to the fact that the preceding DSS-crosslinking to immobilize the antibody on the beads masked most reactive sites (primary amine groups) available. After LC-MS/ms analysis of total protein components obtained using this approach, we found more proteins in SNTA1-specific pulldown complexes from the same lysate with equal protein levels. Here we identified a total of 38 proteins which were filtered for background proteins. Common proteins identified in both the samples like ACTA1, Stress-70 protein, 40S ribosomal protein, Filamin A isoform 2 and CCDC124 were considered to be non-specific proteins. Using this approach we identified 31 SNTA1 binding partners from the MCF-7 cells including motor proteins, actin binding proteins, RNA binding proteins, chaperones and few membrane bound proteins (**Figure 2**). Proteins that fell into the category cytoskeletal proteins included many proteins involved in various cellular processes like cell division, cell motility, cell contractility and organization of organelle transport. The variety of external stimuli trigger signaling cascades implicated in cytoskeletal rearrangement (Lambrechts et al., 2004).

Interestingly, we have identified a very distinct protein repertoire that contributes to breast cancer aggressiveness and metastasis. Some motor proteins identified are involved in protein trafficking towards plasma membrane and also play a major role in crosstalk between tumor microenvironment and cancer cells to elicit cell survival signals and regulate breast cancer cell metastasis (Surcel et al., 2019; Wang et al., 2015; Ouderkirk et al., 2014).

Of the total proteins 28 were found exclusively by crosslinking IP method and 3 proteins (HSPB1, HSP90AA1, TPM3) were found in both groups. Importantly, among the SNTA1-binding partners summarized, we have identified proteins like SNTB1, SNTB2 and Utrophin that had previously been shown to interact with the SNTA1 protein using other strategies, supporting the efficacy of our study (Ahn et al., 1995). Utrophin has been reported previously to interact with alpha-Syntrophin directly via its C-terminal in mouse skeletal muscle cells. These proteins also form a part of Dystrophin Glycoprotein complex to complete a link from the cytoskeleton via the cell membrane and to the extracellular matrix.

## Discussion

In this study, use of Peirce cross-linking CO-IP kit followed by tandem mass spectrometry offered an efficient way of identifying potential SNTA1 binding partners. Crosslinking improves the sensitivity of the method in order to permit the identification of very low-abundance SNTA1 associated proteins which may serve as biomarkers of breast cancer or as targets of novel therapeutics. Using this technique we identified several SNTA1-interacting proteins, as exemplified by the detection of utrophin and syntrophin isoforms like SNTB1 and SNTB2 which were not detected in conventional CO-IP.

Pathway Analysis tool in reactome revealed the pathways exhibited by SNTA1 associated proteins and **Figure 3** shows generalized overview of pathways involved.

**Figure 3:**
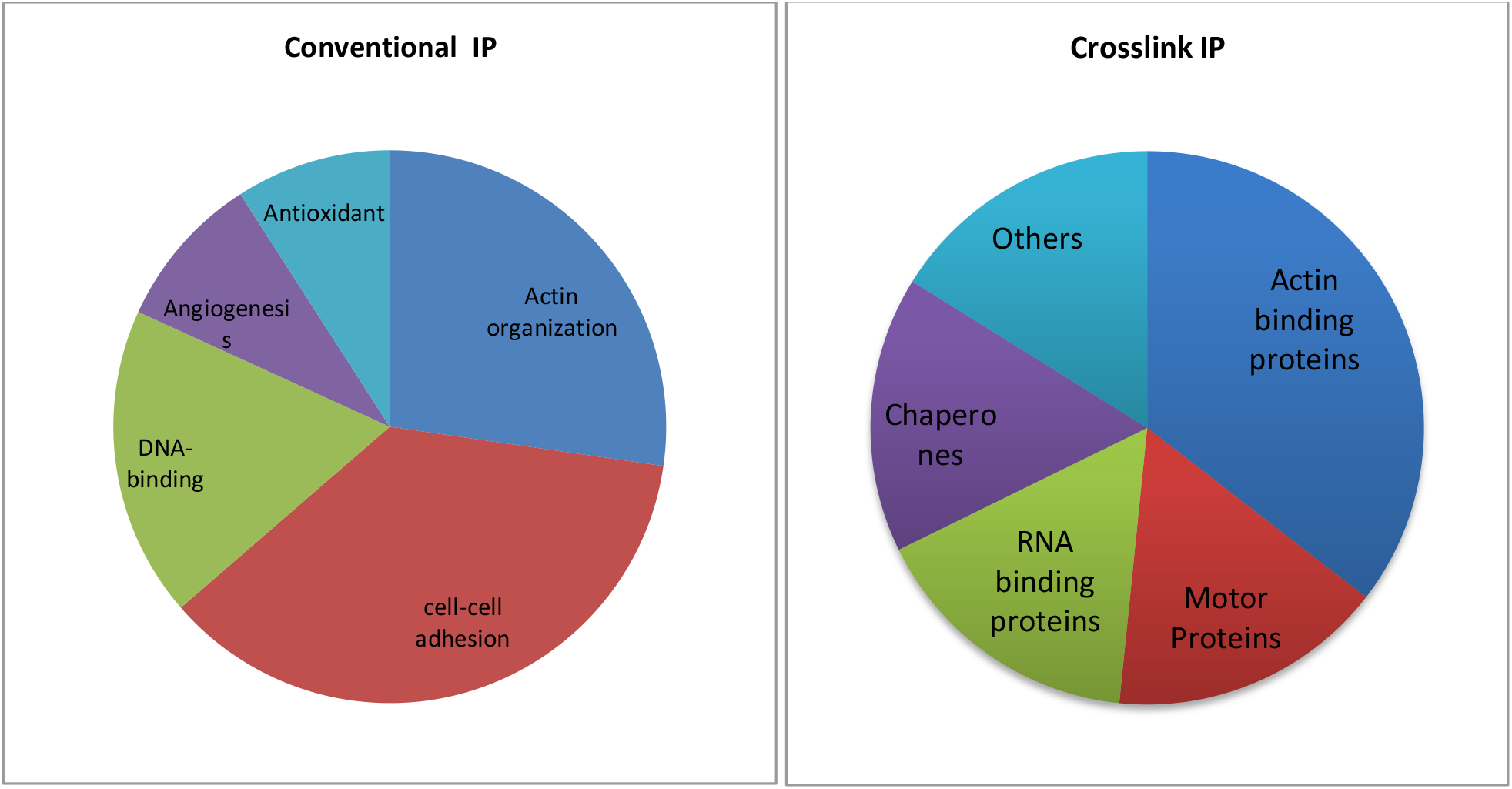
A pie-chart distribution of the proteins identified by conventional and crosslink IP kit into different functional categories.

**Figure 3:**
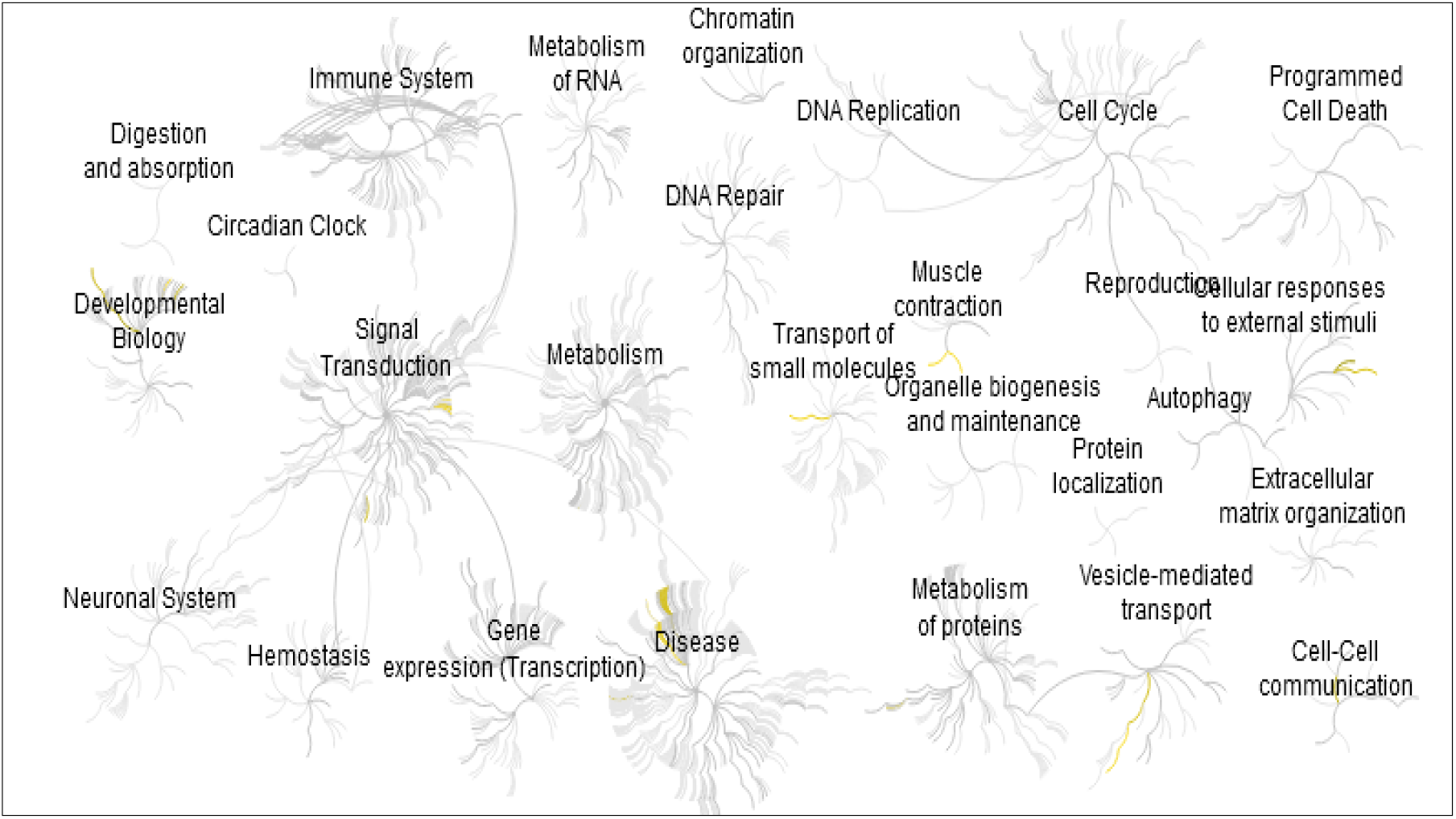
Genome-wide overview of the results of pathway analysis.

Generalized overview of various pathways covering SNTA1 interactome include cell-cell communication, signal transduction, extracellular matrix organization, cell cycle, programmed cell death, muscle contraction, vesicle mediated transport, cellular response to external stimuli and many more. Cell-cell communication and signal transduction regulates migration and growth of neighboring malignant cells.

Even though signaling cascades control all these response mechanisms but a separate group of signal transduction (ID: R-HSA-162582) super pathway in Reactome maps to numerous pathways that are linked to each other in one or more ways and impact cellular proliferation, differentiation, and survival. The best represented pathways also cover some known downstream pathways of SNTA1 signaling, which include Myopathies and Muscle contraction, demonstrating the efficiency of our experiments for determining signal transduction networks. The diversity of proteins identified in this study points to the importance of SNTA1 in the regulation of breast cancer. Furthermore these set of proteins may possess important clinical implications in breast cancer pathology and serve as prognostic/diagnostic biomarkers and therapeutic targets.

## Notes

### Competing Interest Statement

The authors have declared no competing interest.

### Summary of Updates

Revised Authorlist

